# Target DNA-Mediated Plasmonic Coupling and Assembly Kinetics of Gold Nanorods for Label-Free Nucleic Acid Detection

**DOI:** 10.64898/2026.06.16.732610

**Authors:** Shubhangi Sharma, Arun Pratap Singh, Susmita Pradhan, Moksh Goel, Navya Gupta, Satyajit Patra

## Abstract

DNA-programmed assembly of plasmonic nanostructures provides a powerful route to couple molecular recognition with optical signal generation. Here, we report the sequence-specific assembly of DNA-functionalized gold nanorods using a sesame allergen-derived DNA biomarker as a molecular bridge. Target-induced assembly produces concentration-dependent assembly growth, plasmon coupling, and distinct assembly kinetics that are readily monitored by absorption spectroscopy, enabling label-free detection of the target DNA in the nanomolar concentration range. The assembled nanorods further produce strong surface-enhanced Raman scattering (SERS) signals arising from plasmonic coupling within the assemblies, extending detection to the picomolar regime without the use of Raman reporters. Quantitative analysis reveals that both the extent and rate of assembly formation are governed by target DNA concentration. These results establish a direct relationship between molecular recognition, assembly growth, plasmonic coupling, and spectroscopic response, highlighting DNA-programmed gold nanorod assembly as a versatile platform for investigating hybridization-driven plasmonic self-assembly and nucleic acid detection.

**Table of Content:** 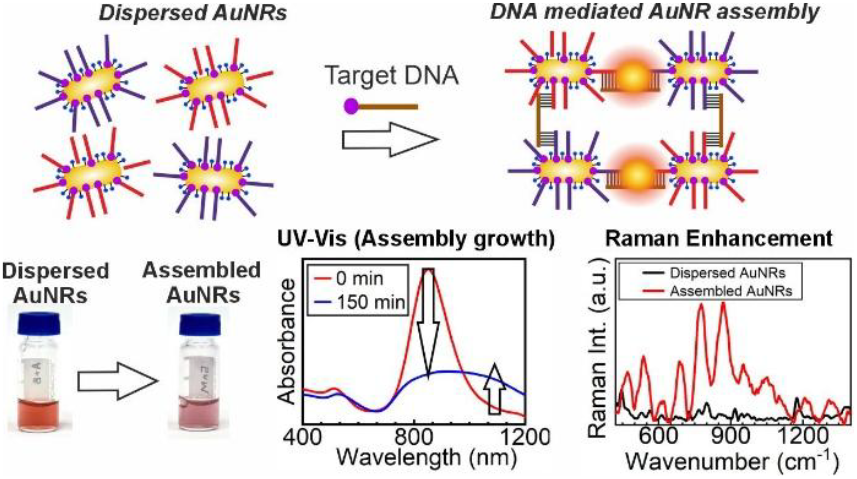

---

Gold nanorods (AuNRs) have attracted considerable interest owing to their anisotropic geometry, tunable localized surface plasmon resonances (LSPRs), and strong electromagnetic field confinement.^1–6^ Unlike spherical nanoparticles, AuNRs support distinct transverse and longitudinal plasmon modes whose energies are highly sensitive to particle aspect ratio and the local dielectric environment.^6–10^ The anisotropic geometry of AuNRs further enables orientation-dependent plasmon coupling,^11,12^ making them attractive building blocks for constructing plasmonic assemblies with tailored optical properties.^3,13^ These characteristics have established AuNRs as versatile platforms for sensing, spectroscopy, imaging, and nanophotonic applications.^1,3,14–16^

The assembly of AuNRs introduces a rich diversity of plasmon coupling modes arising from their anisotropic geometry.^6,11,12^ Unlike spherical nanoparticles, where plasmon coupling is primarily governed by interparticle separation,^17,18^ the optical response of assembled AuNRs depends strongly on both the separation and relative orientation of neighboring nanorods.^6,11,12,19,20^ Consequently, side-by-side and end-to-end assemblies exhibit distinct plasmonic responses arising from different plasmon hybridization modes.^12^ Such orientation-dependent coupling gives rise to unique collective optical properties and intense electromagnetic fields localized within interparticle junctions,^6,11,12^ making assembled AuNR architectures attractive platforms for manipulating light–matter interactions and enhancing spectroscopic signals.^3,21,22^

DNA hybridization provides a powerful and programmable strategy for directing the assembly of plasmonic nanostructures.^13,23–36^ Owing to the high specificity of Watson–Crick base pairing, DNA can function simultaneously as a structural linker and a molecular recognition element, enabling the formation of target-responsive nanoparticle assemblies.^37,38^ DNA-mediated assembly of AuNRs has emerged as a versatile platform for plasmonic sensing, where target molecules trigger nanoparticle assembly through sequence-specific hybridization.^37,39,40^ Previous studies have demonstrated that assembly geometry plays a critical role in determining plasmonic behavior,^6,12,41^ while DNA-directed assembly offers a convenient route for controlling interparticle interactions and collective optical properties.^32,33^

Despite significant advances in DNA-directed AuNR assemblies, most studies have primarily focused on engineering assembly geometry,^13,42–44^ optimizing sensing performance,^27,37^ or characterizing the optical response of the final assembled structures.^6,11^ While previous studies have established the importance of assembly geometry and endpoint optical response,^37,38,45,46^ the influence of target concentration on the growth and evolution of AuNR assemblies remains less understood. In particular, quantitative relationships between molecular recognition, assembly growth, plasmonic coupling, and spectroscopic response have not been systematically established. Addressing these relationships is important for both the rational design of assembly-based biosensors and the broader understanding of collective plasmonic phenomena in anisotropic nanoparticle assemblies.

Herein, we report a target DNA-mediated assembly of DNA-functionalized AuNRs for label-free nucleic acid detection. The target DNA acts as a molecular bridge between two DNA-functionalized AuNR probes containing sequences complementary to different regions of the target, thereby driving the formation of plasmonically coupled assemblies. By systematically varying the target concentration and monitoring the temporal evolution of the optical response, we investigate how molecular recognition influences assembly growth and assembly kinetics. Microscopic characterization reveals progressively larger AuNR assemblies with increasing target concentration, while absorption spectroscopy demonstrates concentration-dependent plasmonic coupling. Together, these results establish a direct relationship between target concentration, assembly growth, plasmonic coupling, and the resulting optical and Raman responses in DNA-mediated AuNR assemblies.

The overall sensing strategy is based on the target DNA-mediated assembly of two populations of DNA-functionalized gold nanorods (AuNRs), as illustrated in **Figure 1A**. To construct the assembly system, citrate-stabilized AuNRs were divided into two portions and independently functionalized with thiolated capture DNA-A and capture DNA-B using a freeze–thaw-assisted conjugation strategy.^47^ Repeated freeze–thaw cycles have been reported to facilitate the loading of thiolated oligonucleotides onto gold surfaces by promoting the formation of stable Au–S bonds and improving surface accessibility of the DNA molecules. The resulting DNA-functionalized AuNRs serve as the fundamental building blocks for the target-directed assembly process. The successful functionalization of the AuNRs with DNA was first evaluated through their colloidal stability under high ionic strength conditions. While citrate-stabilized AuNRs typically undergo aggregation in concentrated salt solutions due to screening of electrostatic repulsion, the DNA-functionalized AuNRs remained stable even in the presence of 500 mM NaCl (**Figure S3, S4**, Supporting Information). The enhanced salt stability indicates the presence of a protective DNA corona that provides both electrostatic and steric stabilization, thereby preventing nonspecific aggregation.^33^ Additional characterization by absorption spectroscopy, zeta potential measurements and thermal reversibility further confirmed the successful loading of DNA on the AuNR surface (**Figures S3–S6**, Supporting Information).

**Figure 1.**
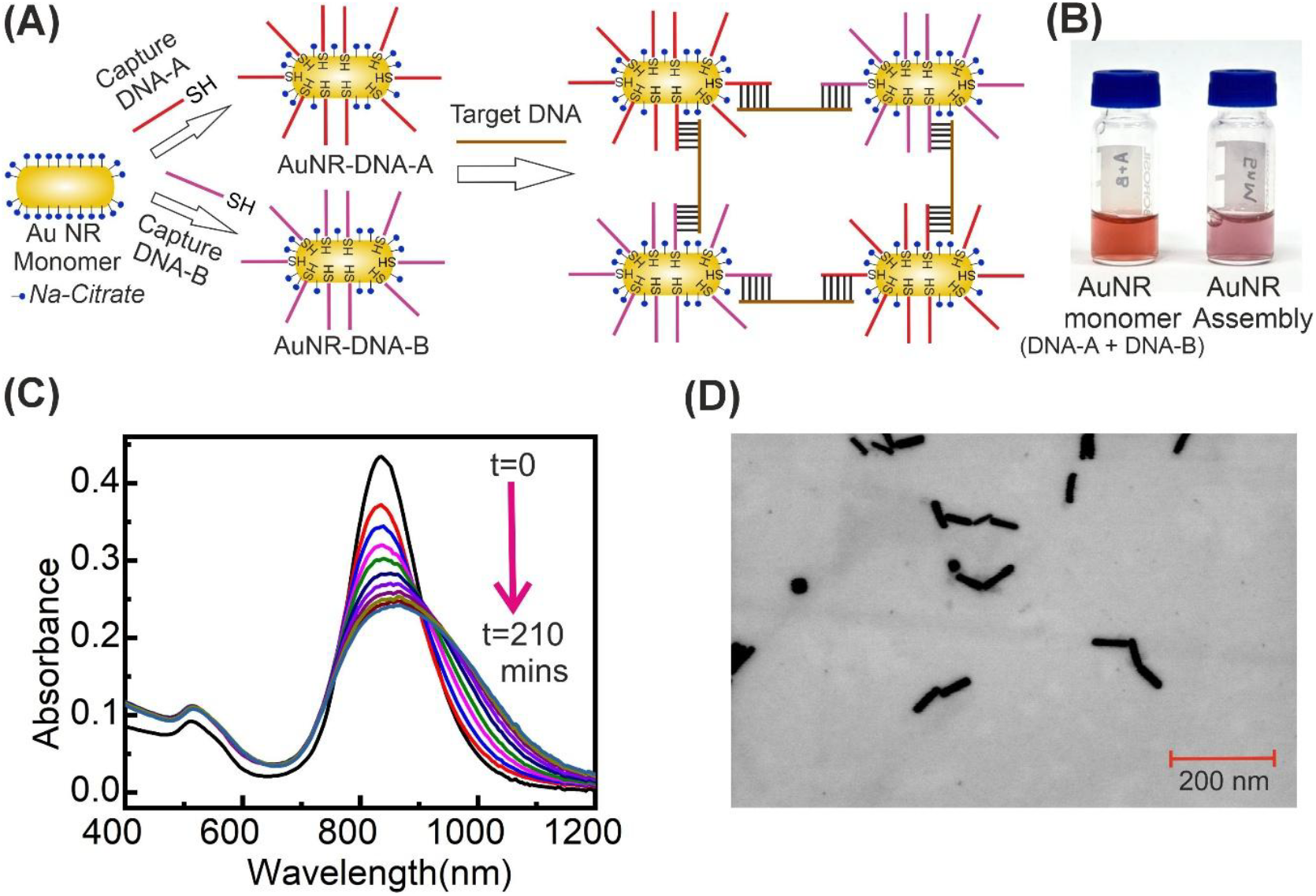
Target mediated coupling of Au nanorods. (A) Schematic presentation of target DNA mediated coupling of AuNRs capped with single strand DNA-A and DNA-B. Target DNA has base segments complementary to the DNA-A and DNA-B and upon hybridization of target DNA with DNA-A and DNA-B attached to the NRs lead to formation of AuNR assembly. (B) Photo of the solution of the DNA capped monomeric AuNR (left) and aggregated AuNR (Right). The color of the solution becomes lighter upon assembly. (C) Plot of absorption spectra as a function of time demonstrating the formation of AuNR assembly. Here the AuNR and target DNA concentrations are kept as 0.5 nM and 5 nM, respectively. As the assembly forms, the absorption at LSPR decreases over time and absorption > 900 nm increases. (D) Scanning tunnelling electron microscopy (STEM) images of the AuNR assembly that exhibits the absorption spectrum in C.

The assembly process exploits the sequence specificity of Watson–Crick base pairing.^32–35^ Capture DNA-A and capture DNA-B are non-complementary to each other and therefore do not induce assembly in the absence of the target strand. However, both capture strands contain regions complementary to distinct segments of the sesame allergen-derived target ssDNA.^48^ Consequently, upon introduction of the target DNA, simultaneous hybridization with both capture strands becomes possible, allowing the target molecule to function as a molecular bridge between neighboring AuNRs. This hybridization-driven bridging interaction promotes the formation of plasmonically coupled AuNR assemblies through sequence-specific molecular recognition.

Formation of the assemblies is accompanied by a visible color change of the AuNR suspension (**Figure 1B**), indicating substantial modification of the plasmonic response of the system. As neighboring nanorods are brought into close proximity through target-mediated hybridization, near-field interactions between their longitudinal plasmon modes give rise to plasmon hybridization within the assembly.^6,20^ Consistent with this interpretation, the absorption spectra exhibit a progressive attenuation of the longitudinal surface plasmon resonance (LSPR) band together with the emergence of broad absorption at longer wavelengths (**Figure 1C**).^11,43^ The appearance of this long-wavelength absorption is characteristic of plasmonically coupled AuNR assemblies and reflects the formation of lower-energy collective plasmon modes arising from interparticle coupling.^6,12,37,41^ Direct evidence for assembly formation was obtained from STEM imaging (**Figure 1D**), which revealed the presence of coupled nanorod structures following addition of the target DNA. The combined optical and microscopic observations confirm that sequence-specific DNA hybridization drives the formation of plasmonically coupled AuNR assemblies.

Having established the target DNA-mediated assembly of DNA-functionalized AuNRs, we next investigated how the assembly process evolves as a function of target DNA concentration. For these experiments, the concentrations of AuNR-A and AuNR-B were kept constant at 0.5 nM, while the concentration of the sesame allergen-derived target DNA was systematically varied. **Figure 2A–D** shows the evolution of the absorption spectra recorded at different incubation times following the addition of 0.25, 5, 10, and 50 nM target DNA, respectively. A strong concentration dependence of the assembly process is immediately evident from the spectral evolution. At the lowest target concentration investigated (0.25 nM), only a modest decrease in the longitudinal surface plasmon resonance (LSPR) band is observed over the course of the experiment, indicating limited assembly formation even after prolonged incubation (**Figure 2E**). In contrast, increasing the target DNA concentration results in progressively larger spectral changes. The attenuation of the LSPR band becomes more pronounced, while the optical density at longer wavelengths increases significantly with time. Such spectral evolution is consistent with enhanced plasmonic coupling arising from the formation of increasingly extensive AuNR assemblies.^1,6,11,41^ The concentration dependence of the optical response reflects the increasing probability of target-mediated bridging interactions between neighboring nanorods as the target DNA concentration increases.

**Figure 2.**
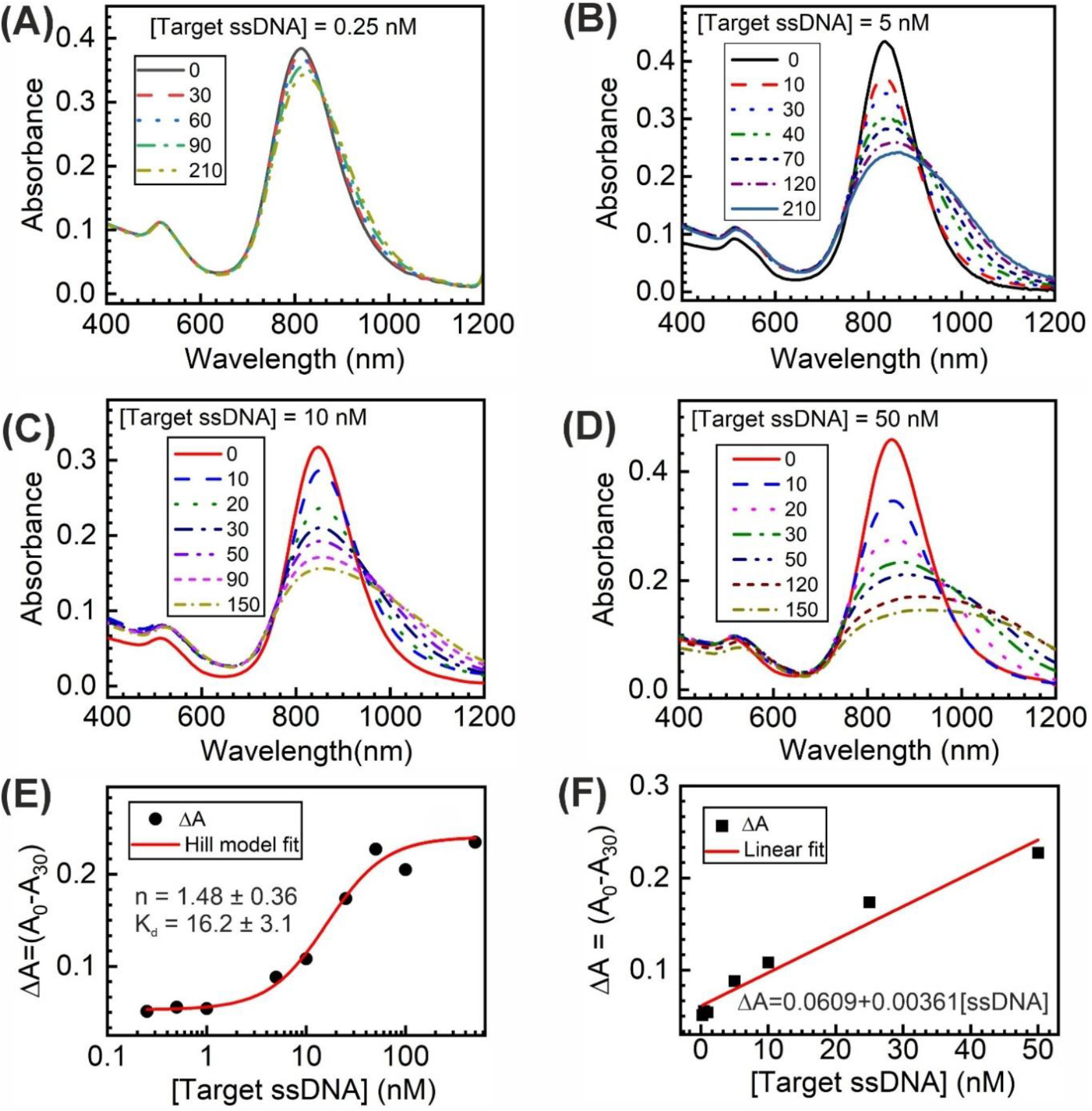
Target concentration-dependent assembly of DNA-functionalized AuNRs monitored by absorption spectroscopy. (A–D) Absorption spectra recorded at different incubation times (in minutes) following the addition of 0.25, 5, 10, and 50 nM target ssDNA, respectively. Progressive attenuation of the longitudinal surface plasmon resonance (LSPR) band together with the emergence of a red-shifted plasmon-coupled band indicates target-mediated nanorod assembly and enhanced plasmonic coupling.^1,6,11,41^ (E) Dependence of the absorbance change (ΔA = A_0_ − A_30_) on target ssDNA concentration. The solid line represents a Hill model fit, yielding an apparent dissociation constant (K_d_) of 16.2 ± 3.1 nM and a Hill coefficient (n) of 1.48 ± 0.36. (F) Linear calibration plot obtained in the low-concentration regime, demonstrating quantitative detection of target ssDNA.

In addition to the extent of assembly, the temporal evolution of the spectra also exhibits a pronounced concentration dependence. At low target concentrations, the optical response develops gradually over several hours, whereas substantially faster spectral changes are observed at higher target concentrations (**Figure 2A-D** and **Figure S7**). The acceleration of the assembly process with increasing target concentration suggests that the availability of target strands strongly influences the formation of plasmonically coupled AuNR assemblies. Furthermore, careful inspection of the spectral evolution reveals that the changes are not strictly self-similar across the concentration range investigated. In particular, systematic variations in the spectral profiles and isosbestic behavior are observed as the target concentration increases (**Figure 2A-D** and **Figure S7**). These observations suggest that the spectral evolution cannot be explained solely by a uniform increase in assembly rate and may instead reflect differences in the size distribution and structural complexity of the assemblies formed at different target concentrations. The interpretation of the absorption data is further supported by STEM characterization (**Figure S8**, Supporting Information), which reveals a progressive increase in the size and complexity of the nanorod assemblies with increasing target DNA concentration. The structural evolution observed by electron microscopy is consistent with the enhanced plasmonic coupling inferred from the absorption spectra. Importantly, the structural evolution observed by STEM correlates closely with the concentration-dependent optical response. At low target concentrations, only small assemblies and short oligomers are observed, resulting in relatively modest changes in the LSPR band. In contrast, higher target concentrations produce larger and more complex assemblies, which exhibit substantially stronger plasmonic coupling and more pronounced long-wavelength absorption.^20^ These observations indicate that the optical response is governed not simply by assembly formation, but by the progressive growth of plasmonically coupled AuNR structures as the concentration of the bridging DNA strand increases.

To quantify the concentration dependence of the assembly response, the absorbance change, ΔA = A_0_ – A_30_, was plotted as a function of target DNA concentration (**Figure 2E**). The response increases sigmoidally with increasing target concentration before approaching saturation at higher concentrations. The data were fitted using the Hill equation to provide an empirical description of the concentration dependence of the assembly process. The resulting Hill coefficient of 1.48 ± 0.36 suggests a positively cooperative assembly response, although the complexity of the assembly process precludes rigorous interpretation in terms of a simple binding equilibrium. The concentration-dependent optical response further enables quantitative detection of the target DNA. As shown in **Figure 2F**, a linear relationship is obtained over the low-concentration regime, allowing construction of a calibration curve for target DNA detection. Using the 3σ/s criterion, where σ is the standard deviation of the blank signal and s is the slope of the calibration plot, the limit of detection was determined to be 1.7 nM. The observed sensitivity demonstrates that the concentration-dependent plasmonic response associated with target-induced AuNR assembly can be exploited for label-free nucleic acid detection using a simple absorption assay.

To gain further insight into the assembly process, the temporal evolution of the optical response was analyzed quantitatively. **Figure 3A** shows the change in absorbance as a function of time for different target DNA concentrations. For all concentrations investigated, the absorbance decreases progressively with time before approaching a plateau value, consistent with the gradual formation of plasmonically coupled AuNR assemblies. The kinetic traces are well described by a single-exponential decay function, allowing extraction of an apparent rate constant (k_app_) that characterizes the overall rate of assembly formation. A pronounced concentration dependence of the assembly kinetics is observed. At low target concentrations, the absorbance evolves slowly and requires a significantly longer time to reach equilibrium. In contrast, the decay becomes progressively faster as the target DNA concentration increases. This behavior is reflected in the extracted k_app_ values, which increase systematically with increasing target concentration (**Figure 3B**). The observed acceleration of the assembly process indicates that the rate of nanorod assembly is strongly influenced by the availability of target DNA molecules capable of simultaneously hybridizing with capture DNA-A and capture DNA-B.

**Figure 3.**
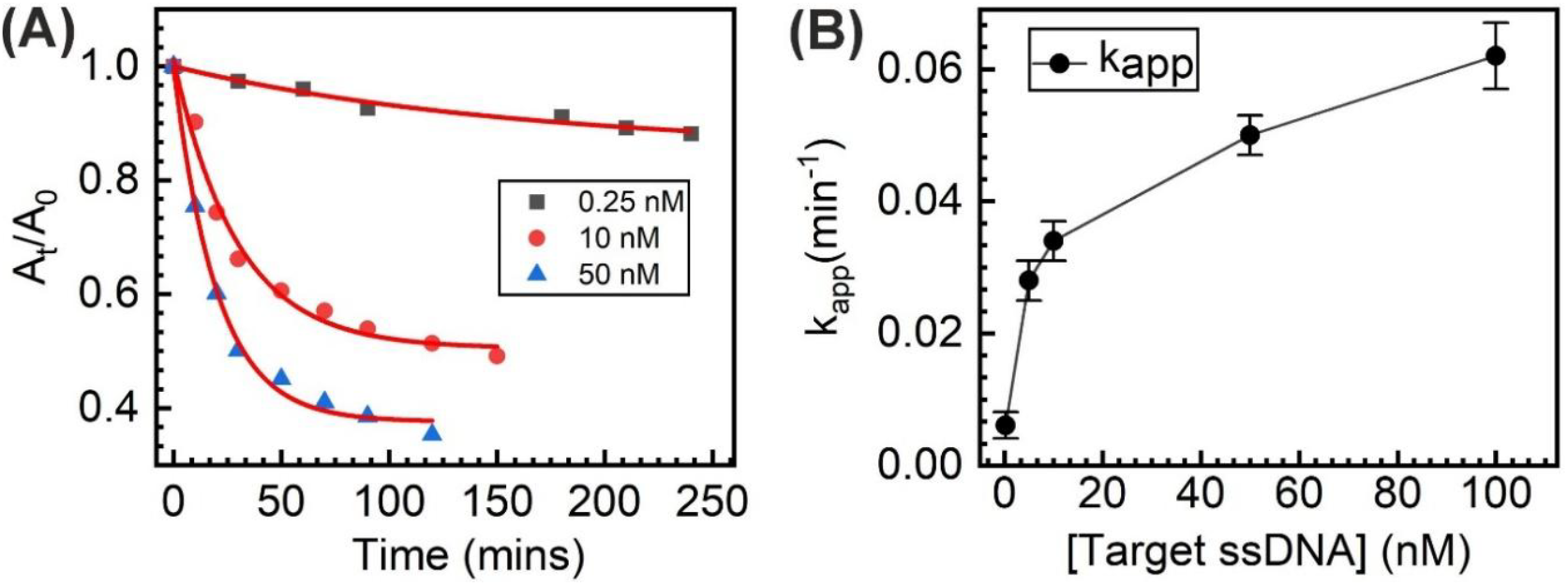
Concentration-dependent assembly kinetics of DNA-functionalized gold nanorods mediated by the sesame-allergen-derived target DNA. (A) Time-dependent changes in the normalized longitudinal plasmon absorbance (A_t_/A_0_) following the addition of 0.25 nM (black squares), 10 nM (red circles), and 50 nM (blue triangles) target DNA. Solid lines represent exponential fits used to determine the apparent assembly rate constants (k_app_). (B) Dependence of the apparent assembly rate constant (k_app_) on target DNA concentration. Error bars represent the fitting uncertainties obtained from the exponential fits. The increase in k_app_ with increasing target DNA concentration indicates accelerated target-mediated nanorod assembly.

The concentration dependence of the assembly kinetics can be understood in terms of the probability of productive target-mediated bridging events. At low target concentrations, encounters between AuNR-A and AuNR-B probes frequently occur without successful target-mediated bridging, resulting in slow assembly growth and prolonged equilibration times. Increasing the target concentration increases the likelihood that a nanorod encounter leads to simultaneous hybridization with both capture strands, thereby accelerating assembly formation. Consequently, the assembly process evolves on progressively shorter timescales as the concentration of the bridging DNA strand increases. It should be noted that the extracted rate constants do not correspond to a single microscopic step in the assembly process. Instead, they reflect the combined contributions of target hybridization, nanorod bridging, and assembly growth that collectively determine the temporal evolution of the absorbance. Nevertheless, the monotonic increase of k_app_ with target concentration clearly demonstrates that the kinetics of AuNR assembly can be effectively modulated through the concentration of the bridging DNA strand. The ability to control both the extent and rate of assembly formation through target concentration provides valuable insight into the dynamic behavior of hybridization-driven plasmonic nanostructure assembly.

Having established the concentration-dependent formation and kinetics of target-induced AuNR assemblies, we next investigated whether the assembled structures could generate plasmonic hotspots capable of enhancing the Raman signal of the DNA. **Figure 4A** compares the Raman spectra of a mixture of AuNR-A and AuNR-B in the absence and presence of the target DNA. In the absence of target DNA, only weak Raman features are observed due to the lack of significant electromagnetic enhancement. Upon addition of the target DNA, however, a substantial increase in Raman intensity is observed across the spectrum. The enhancement coincides with the formation of AuNR assemblies and is consistent with the generation of localized electromagnetic hotspots within the interparticle junctions of the coupled nanorods.^49–51^ The strong confinement of the electromagnetic field within these nanogaps is expected to amplify the Raman scattering cross section of nearby DNA molecules, resulting in the substantial increase in Raman intensity observed after assembly formation.^49,50^

**Figure 4.**
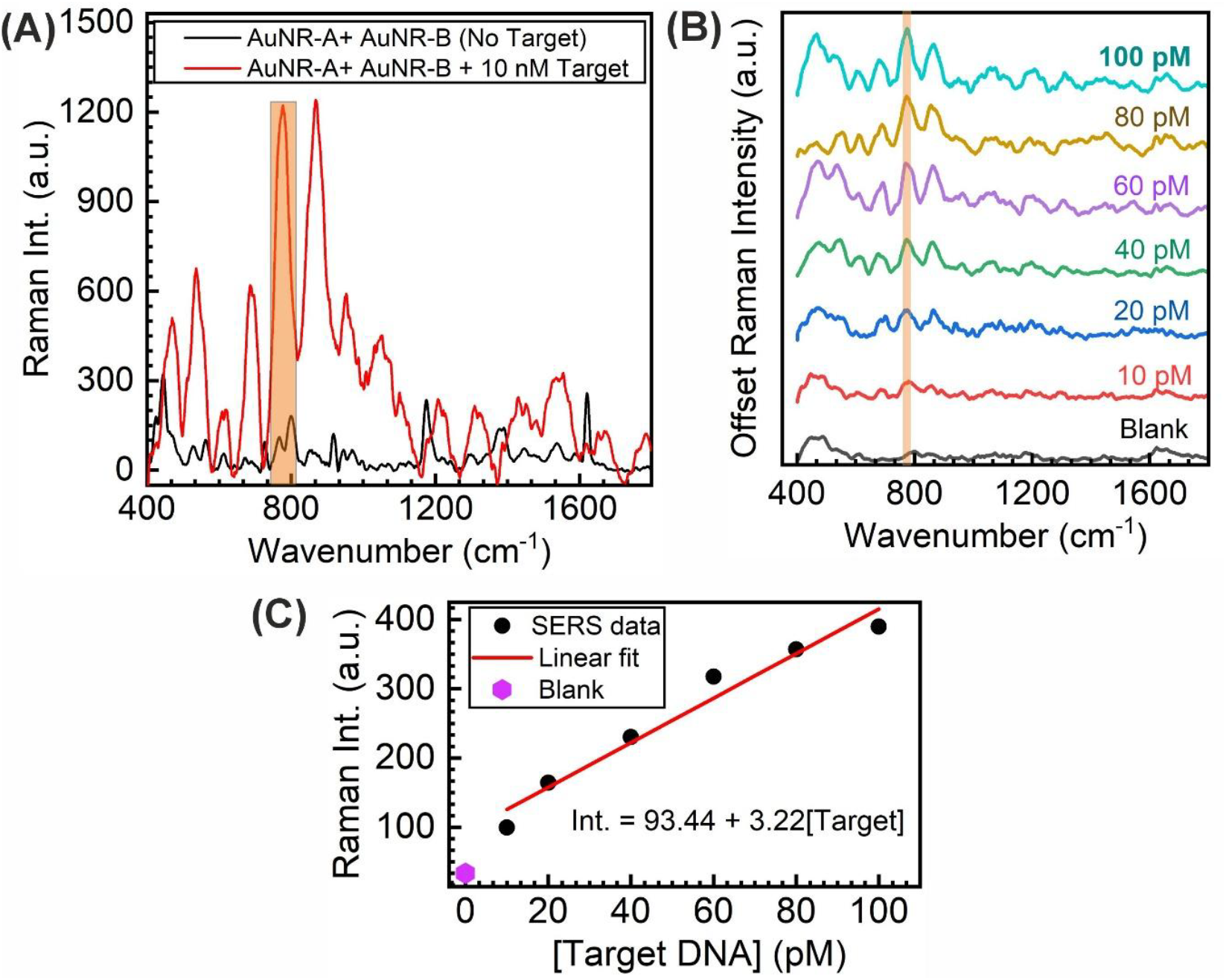
Target-induced plasmonic hotspot formation and SERS detection of the sesame-allergen-derived DNA target. (A) SERS spectra of the mixed AuNR-DNA-A and AuNR-DNA-B system in the absence and presence of 10 nM target DNA, demonstrating substantial signal enhancement upon target-mediated assembly. (B) Concentration-dependent SERS spectra recorded in the presence of increasing concentrations of target DNA (10–100 pM). The highlighted region corresponds to the characteristic Raman band at 773 cm^−1^ used for quantitative analysis. Spectra are vertically offset for clarity. (C) Dependence of the Raman intensity of the 773 cm^−1^ band on target DNA concentration. The solid line represents a linear fit to the experimental data, demonstrating concentration-dependent enhancement of the SERS response.

Several characteristic Raman bands become clearly visible following assembly formation (**Figure 4A**). The Raman band at approximately 773 cm^−1^ was selected for quantitative analysis because it exhibits one of the highest signal-to-noise ratios among the observed vibrational features and displays a systematic dependence on target concentration. The assignment is further supported by comparison with the Raman spectra of individual nucleotide monophosphates (**Figure S9**), where adenine-containing species exhibit prominent Raman features in the 770–800 cm^−1^ region. The appearance and amplification of these vibrational signatures confirm that the assembled AuNR structures provide sufficient electromagnetic enhancement to enable direct detection of DNA without the use of externally added Raman reporter molecules. Notably, no detectable Raman signal could be observed from the target DNA alone at concentrations as high as 20 μM under identical experimental conditions (**Figure S10**, Supporting Information). In contrast, strong Raman signals are readily observed following target-induced assembly of the AuNRs at nanomolar DNA concentrations, highlighting the substantial signal amplification generated by the assembly-induced plasmonic hotspots.

To evaluate the analytical performance of the assembly-generated hotspots, Raman spectra were recorded at different target DNA concentrations (**Figure 4B**). The intensity of the characteristic Raman bands increases progressively with increasing target concentration, consistent with the formation of a larger population of plasmonically active assemblies. For quantitative analysis, the intensity of the 773 cm^−1^ band was extracted and plotted as a function of target DNA concentration (**Figure 4C**). A monotonic increase in Raman intensity is observed with increasing target concentration, enabling construction of a calibration curve for SERS-based detection. Using the same 3σ/s criterion employed for the spectrophotometric assay, the limit of detection for the SERS assay was determined to be 2 pM. The substantially lower detection limit compared to the spectrophotometric assay (1.7 nM) highlights the enhanced sensitivity of the plasmonically coupled AuNR assemblies.

The enhanced sensitivity of the Raman response relative to the spectrophotometric assay arises from the strong localization of the electromagnetic field within the interparticle gaps of the assembled nanorods. As a result, target-induced assembly not only produces measurable plasmonic coupling in the absorption spectra but also generates regions of enhanced electromagnetic fields that substantially amplify the vibrational signatures of the DNA. The strong electromagnetic enhancement generated within the plasmonically coupled AuNR assemblies enables reporter-free Raman detection of the target DNA in the picomolar concentration regime, demonstrating the potential of the assembled AuNR platform for sensitive nucleic acid analysis.

In summary, we have demonstrated the sequence-specific assembly of DNA-functionalized gold nanorods using a sesame allergen-derived target DNA as a molecular bridge. Hybridization-driven assembly produced pronounced concentration-dependent plasmonic responses, enabling direct monitoring of both the extent and kinetics of assembly formation through absorption spectroscopy. Quantitative kinetic analysis revealed that the rate of assembly formation increases systematically with target DNA concentration, highlighting the dynamic nature of the assembly process. The resulting AuNR assemblies further generated plasmonically coupled nanostructures capable of producing strong SERS enhancement, enabling reporter-free detection of the target DNA with significantly improved sensitivity compared to the spectrophotometric assay. The combination of target-mediated assembly, concentration-dependent plasmonic coupling, and Raman enhancement establishes a direct relationship between molecular recognition, assembly growth, and spectroscopic response. Beyond its analytical utility, this study provides insight into how target concentration governs the growth and plasmonic response of anisotropic nanoparticle assemblies, contributing to a broader understanding of hybridization-driven plasmonic self-assembly and its application in nucleic acid detection.

## Supporting information

Data is enclosed as supplementary files

## Acknowledgements

This study is supported by the Start-up Research Grant (SERB/SRG/2022/000341) from the Anusandhan National Research Foundation (ANRF), Government of India. Financial support from the Student Program for Advancing Research, KnowLedge, and Entrepreneurship (SPARKLE) project of BITS Pilani is also gratefully acknowledged. Additional support is provided through the Research Initiation Grant (RIG) and Additional Competitive Research Grant (ACRG) of BITS Pilani. S. Pradhan, S. Sharma, and A. P. S. acknowledge Institute Fellowships from BITS Pilani. S. Sharma also acknowledges the Junior Research Fellowship (JRF) received from ANRF during November 2022–September 2024.

