## Supplementary material for "Target DNA-Mediated Plasmonic Coupling and Assembly Kinetics of Gold Nanorods for Label-Free Nucleic Acid Detection": Data is enclosed as supplementary files

### Contents:

**1.** Materials and methods: Chemicals, Sample Preparation and Experimental Methods

**2.** Normalized absorption spectrum of Au NRs before and after citrate capping (**Figure S1**)

**3.** Fourier-transform infrared (FTIR) spectrum of citrate and CTAB capped Au NRs (**Figure S2**).

**4.** Absorption spectra of DNA functionalised AuNRs after purification (**Figure S3**)

**5.** Salt-induced aggregation of citrate functionalized Au NRs monitored by absorption spectroscopy (**Figure S4**)

**6.** Zeta Potential measurements during AuNR surface modification and DNA functionalization (**Figure S5**)

**7.** Thermal reversibility of DNA-mediated AuNR assemblies (**Figure S6**)

**8.** UV–Vis spectral evolution of target-induced AuNR assemblies at 100 nM and 500 nM target DNA concentrations (**Figure S7**)

**9.** STEM images of target DNA-mediated AuNR assemblies formed at different target DNA concentrations (**Figure S8**)

**10.** Normal Raman spectra of nucleotide monophosphates (**Figure S9**)

**11.** Normal Raman spectrum of the target DNA recorded at a concentration of 20 µM under identical experimental conditions used for the SERS measurements (**Figure S10**)

### 1. Materials and Methods:

#### 1.1. Chemicals and Sample Preparation:

All the DNA oligonucleotides are purchased from Sigma-Aldrich (Merck). The sequences of these oligonucleotides are the following.

**Capture DNA-A (Cap A):** 5′-AAA-GAT-AGC-CCA-AAA-ATT-CAA-AGG-ATT-TTT-C-6Thio-3′

**Capture DNA-B (Cap B):** 5′-ThioC6-TT-TTA-GAA-TTT-GAC-TCC-GTA-CCA-CTG-AAG-3′

**Target ssDNA:** 5′-TCC-TTT-GAA-TTT-TTG-GGC-TAT-CTT-TCA-AGC-GTG-CGA-ATG-AAC-CCT-TCA-GTG-GTA-CGG-AGT-CAA-ATT-CT-3′

The Cap A and Cap B DNA were hybridized by annealing at 90 °C for 5 min, and then gradually cooled to room temperature in the PCR thermocycler (T-100, Bio-Rad). Chloroauric acid trihydrate ($HAuCl_4 \cdot 3H_2O$), cetyltrimethylammonium bromide (CTAB), L-ascorbic acid, silver nitrate ($AgNO_3$), Hydrochloric acid (HCl), Sodium Borohydride ($NaBH_4$), Sodium Chloride (NaCl, ACS reagent, ≥ 99.0%), Tween-20, Trisodium Citrate (TSC), Polystyrene sulfonate sodium salt (Na-PSS) are also procured from Sigma-Aldrich (Merck).

#### 1.2. Preparation of Au Nanorods (NRs):

Au NRs were prepared using the seed-mediated growth approach.[1,2] First, Au nanoseeds were synthesized in an aqueous solution by reducing $HAuCl_4{\cdot}3H_2O$ (25 µL of 50 mM) in CTAB (4.7mL, 0.1 M) using a freshly prepared solution of $NaBH_4$ (300 µL, 10 mM) under vigorous stirring at 30 °C for 5 minutes. The solution colour changed from yellow to brown within a few seconds, confirming the formation of Au nanoseeds. Next, the Au nanoseed solution was aged at 30 °C for 1 hour. After that, the nanorod growth solution was prepared by adding 190 µL of 1 M HCl and 100 µL of 50 mM $HAuCl_4$ to 10 mL of CTAB (0.1 M) solution, and gently stirring for 5 minutes until the solution became homogeneous. The pH of the solution was 1.5. Then, 120 µL of 10 mM $AgNO_3$ was added under mild stirring. After that, 100 µL of 100 mM Ascorbic acid was added, turning the solution from yellow to colourless, indicating the reduction of $Au^{3+}$ to $Au^{+}$. Finally, 24 µL of the aged seeds was added, and the solution was left undisturbed overnight. The solution turned deep red, indicating the formation of Au NRs. To remove excess CTAB and unreacted chemicals, the solution was centrifuged at 9500 rpm for 10 minutes and stored in 10 mM CTAB for future use.

#### 1.3. Ligand Exchange and Surface Modification of Au NRs:

##### 1.3.1. Preparation of Citrate-stabilized Au NRs:

CTAB-capped Au NRs (1 mL) were diluted with 1 mL of Milli-Q water and centrifuged at 9500 rpm for 20 min to remove excess CTAB. The resulting pellet was redispersed in 1 mM CTAB, and the purification process was repeated twice. The partially CTAB-depleted Au NRs were subsequently redispersed in 1 mL of 0.20 wt% sodium polystyrene sulfonate (Na-PSS) and subjected to four cycles of centrifugation and redispersion to achieve complete ligand exchange.[3] The resulting PSS-coated Au NRs were stable at room temperature for several weeks. Finally, the PSS-coated Au NRs were transferred into 5 mM trisodium citrate (TSC) through three successive centrifugation and redispersion cycles.[3] Excess citrate was removed by centrifugation, and the citrate-stabilized Au NRs were finally dispersed in Milli-Q water.

Successful surface modification was confirmed by UV–Vis absorption and FTIR spectroscopy (**Figure S1 and S2**). Following ligand exchange, a slight blue shift of the longitudinal plasmon resonance band was observed. This shift is attributed to the replacement of the CTAB bilayer with citrate, which possesses a lower local refractive index surrounding the AuNR surface.

#### 1.3.2. Preparation of DNA Functionalization of Au NRs:

For DNA functionalization of AuNRs, two sequences of thiol-modified oligonucleotides were used, named Cap A and Cap B. Two aliquots of citrate-stabilised Au NRs (NR@Cit) were prepared, and 1 µM of Cap A and Cap B were added separately with 0.1wt% of Tween 20 and 0.01% of SDS. The sample was frozen overnight and thawed the next day to promote DNA adsorption on the nanoparticle surface.[4] The solutions were centrifuged three times to remove excess DNA, Tween 20, and SDS; finally, 500 mM NaCl was added, and the particles were mixed. Next, Different concentrations of target ssDNA were added to initiate DNA hybridization, and the reaction was monitored by absorption spectroscopy.

#### 1.4. Zeta Potential measurements:

Zeta Potential measurements were conducted on a Litesizer 500 (Anton Paar) using a univette low-volume cuvette (60 µL). The measurements were conducted in aqueous solution at room temperature ($25^0$C).

#### 1.5. Absorption spectroscopy measurements:

Absorption spectra of AuNRs were recorded on a UV-Vis-NIR spectrophotometer (Lambda 1050+, PerkinElmer), with a cuvette volume of 1 mL and a path length of 0.2 cm (Starna). The optical density of AuNRs was kept at 0.5. The absorption spectra were recorded in water at room temperature ($25^0$C) for all samples.

#### 1.6. Fourier-transform infrared spectroscopy measurements:

The FTIR spectra were acquired using a Shimadzu IRAffinity-1S spectrometer. CTAB-capped and citrate-capped nanorods were purified by repeated centrifugation to remove excess free ligands. The purified samples were drop-cast multiple times onto IR-grade KBr and dried overnight to ensure the complete removal of residual moisture. Pelleted NRs were prepared and analysed in the respective range. This procedure minimized the broad O-H stretching vibration of water (3200-3400 $cm^{-1}$). The FTIR spectrum of CTAB-capped Au NRs exhibited characteristic alkyl C-H stretching vibrations at 2920 and 2850 $cm^{-1}$, confirming the presence of CTAB bilayer on the nanorod surface. After ligand exchange, these bands completely disappear in the citrate-capped Au NRs, indicating the effective removal of CTAB and Au NRs displayed prominent carboxylate stretching vibrations in the 1600-1400 $cm^{-1}$ region, corresponding to asymmetric and stretching modes of $COO^-$ groups. The disappearance of the

CTAB-associated ν($CH_2$) bands together with the appearance of $COO^-$ stretching vibrations confirms the successful replacement of CTAB by citrate on the Au NR surface.

**1.7. Scanning Transmission Electron Microscopy (STEM) Imaging experiments:**

STEM images were acquired using a field-emission scanning electron microscope (FE-SEM, Apreo LoVac, FEI) operated in STEM mode. For sample preparation, 10 µL of each sample was deposited onto a 300-mesh copper grid and dried in an oven at 40 °C prior to imaging.

**1.8. Raman Measurements**:

Raman and surface-enhanced Raman scattering (SERS) measurements were performed using a confocal Raman microscope (LabRAM HR Evolution, Horiba) at the Central Instrumentation Facility (CIF), BITS Pilani. The samples were excited using a 532 nm laser with a laser power of 100 µW (0.1% of the maximum laser power). All measurements were carried out in the liquid state with an acquisition time of 20 s and 4 accumulations. For SERS measurements, the concentration of DNA-functionalized Au NRs was maintained at 0.5 nM. The Raman spectra of nucleotide monophosphates and target DNA were recorded under identical instrumental conditions.

**2. Normalized absorption spectrum of Au NRs before and after citrate capping**

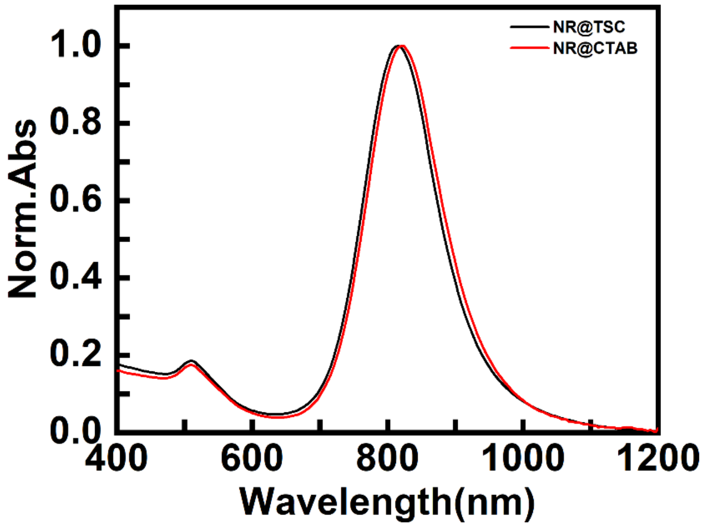


**Figure S1:** Normalised absorption spectra of AuNRs, before (red line) and after citrate capping (black line)

**3. Fourier-transform infrared (FTIR) spectrum of citrate and CTAB capped Au NRs**

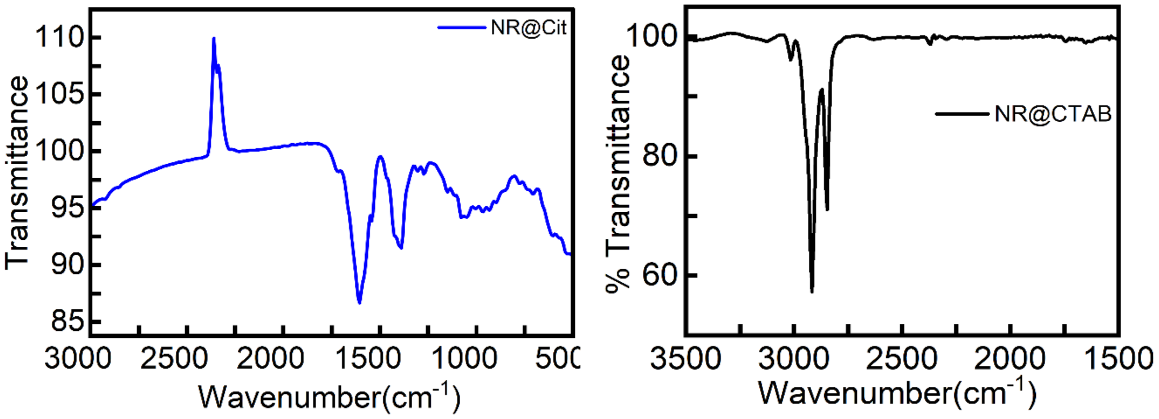


**Figure S2:** FTIR characterisation of citrate-capped AuNRs. The NR@CTAB spectrum exhibits characteristic of alkyl C-H stretching vibrations at 2920 and 2850 cm-1, confirming the presence of the CTAB bilayer on the nanorod surface, which is completely absent in citrate-capped AuNRs. The stretching vibrations of carboxylate groups in the 1600-1400 $cm^{-1}$($COO^{-}$) confirm the ligand exchange.

**4. Absorption spectra of DNA functionalised AuNRs after purification (Figure S3)**

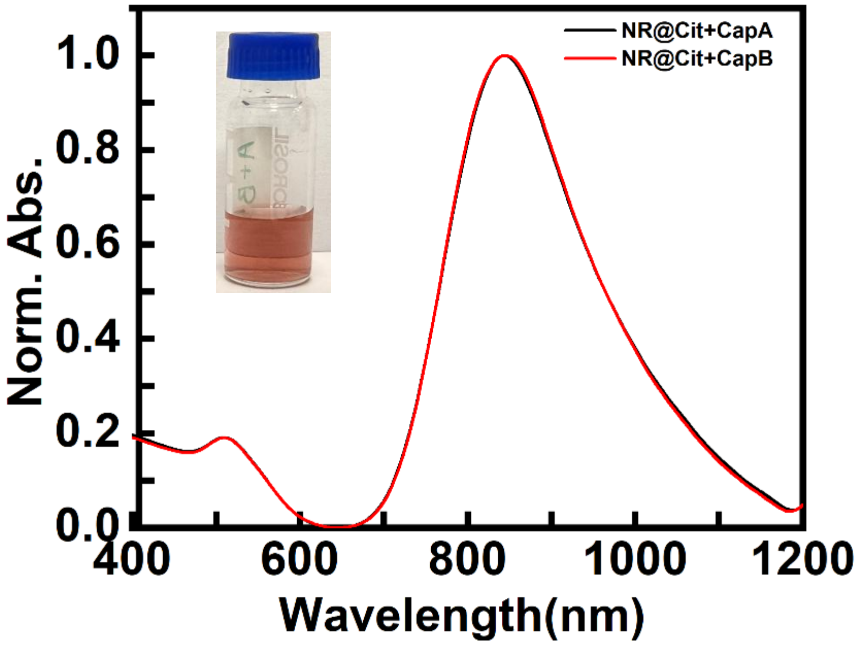


**Figure S3:** Normalised absorption spectra of ssDNA functionalised Au NRs after 3-times centrifugation in 500 mM NaCl. Inset shows the photograph of the DNA capped Au NR solution in 500 mM NaCl. The retention of the color of the Au NR indicates that the NRs remain stable even at such high concentrations of salt.

**5. Salt-induced aggregation of citrate functionalized Au NRs**

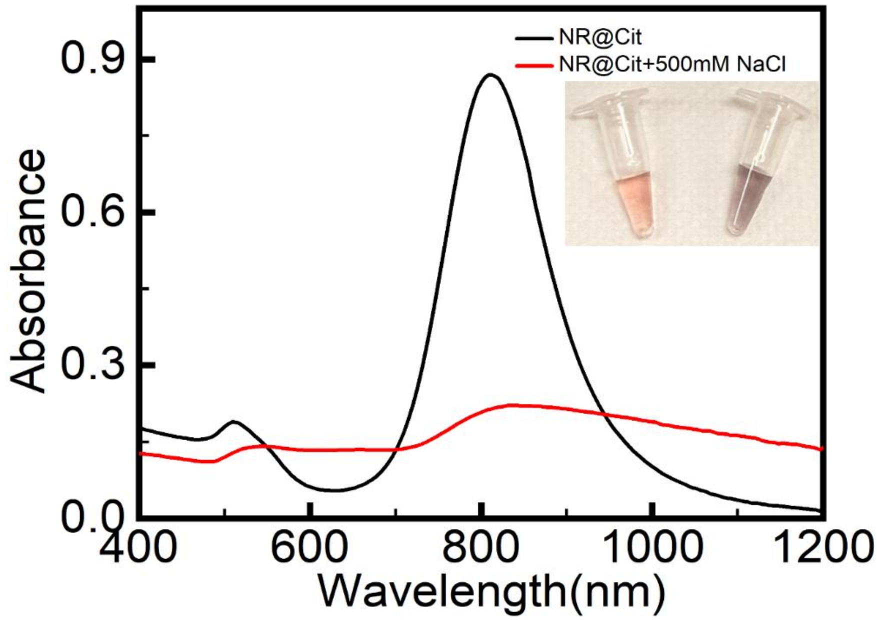


**Figure S4:** Absorption spectra of citrate functionalised AuNRs before and after addition of 500mM NaCl. The high ionic strength screens the electrostatic repulsion. The inset images show the colour change in the colloidal suspension.

### 6. Zeta Potential measurements

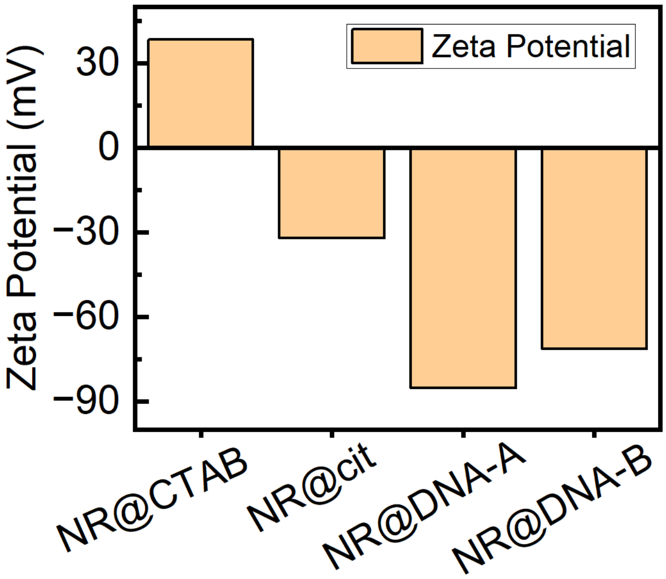


**Figure S5:** Zeta potential measurements of CTAB-capped AuNRs (NR@CTAB), citrate-capped AuNRs (NR@Cit), and DNA-functionalized AuNRs (NR@DNA-A and NR@DNA-B). The positive zeta potential of NR@CTAB (+38.4 mV) arises from the cationic CTAB bilayer. Ligand exchange with citrate reverses the surface charge to −32 mV, confirming successful replacement of CTAB by citrate. Subsequent functionalization with DNA further shifts the zeta potential to −85 mV (NR@DNA-A) and −71 mV (NR@DNA-B), consistent with the presence of negatively charged phosphate groups on the DNA backbone.

**7. Thermal Reversibility:**

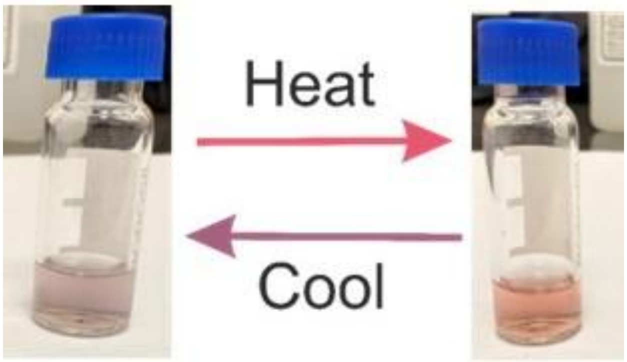


Au NRs assembly Dispersed AuNRs

**Figure S6**: **Thermal reversibility of target DNA-mediated AuNR assemblies**. Photographs showing the optical response of the AuNR assembly upon heating and subsequent cooling. Heating induces disassembly of the AuNR assembly through thermal disruption of DNA hybridization, resulting in a decrease in plasmonic coupling and a corresponding color change. Upon cooling, the complementary DNA strands rehybridize, leading to reassembly of the AuNRs and recovery of the original color. The reversible optical response confirms that the assembly process is governed by DNA hybridization.

**8. UV–Vis spectral evolution of Au NR assemblies higher target DNA concentrations**

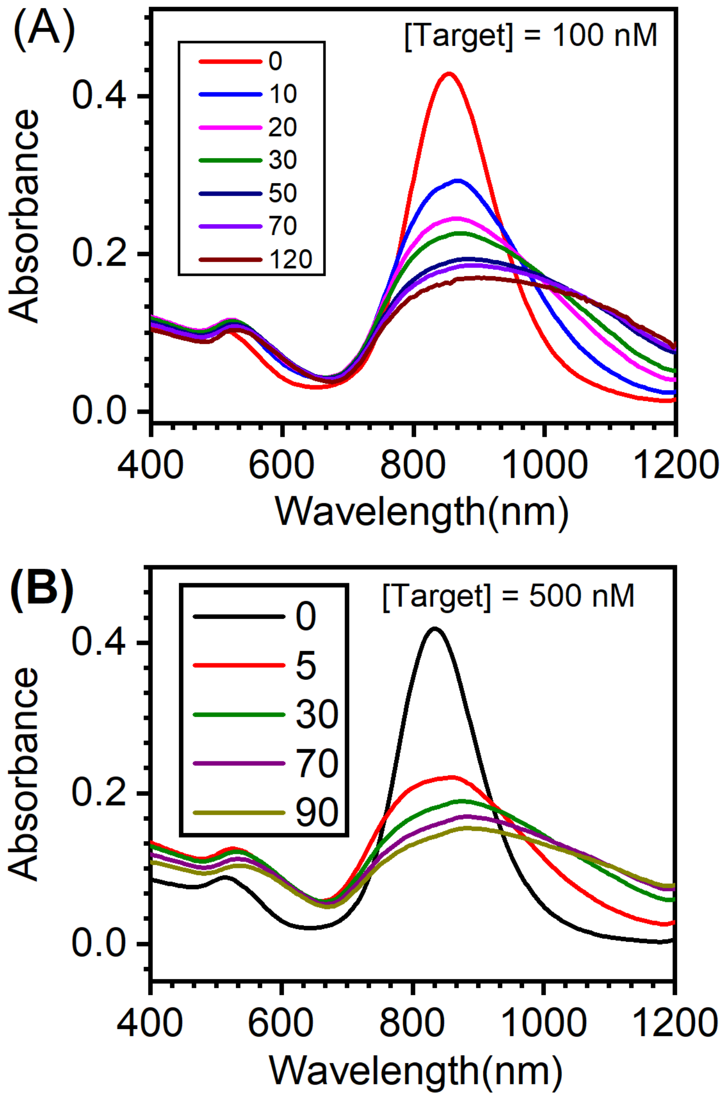


**Figure S7**: **Evolution of the absorption spectra as a function of time for the target-induced AuNR assemblies at high target DNA concentrations.** Absorption spectra of DNA-functionalized Au NRs recorded at different incubation times (in minutes) following the addition of (A) 100 nM and (B) 500 nM target DNA. The concentrations of Au NR-A and Au NR-B were maintained at 0.5 nM. With increasing incubation time, the longitudinal surface plasmon resonance (LSPR) band progressively decreases in intensity, while the optical density at longer wavelengths increases, consistent with the continued growth of target DNA-mediated AuNR assemblies.

**9. Scanning Transmission Electron Microscopy (STEM) Imaging:**

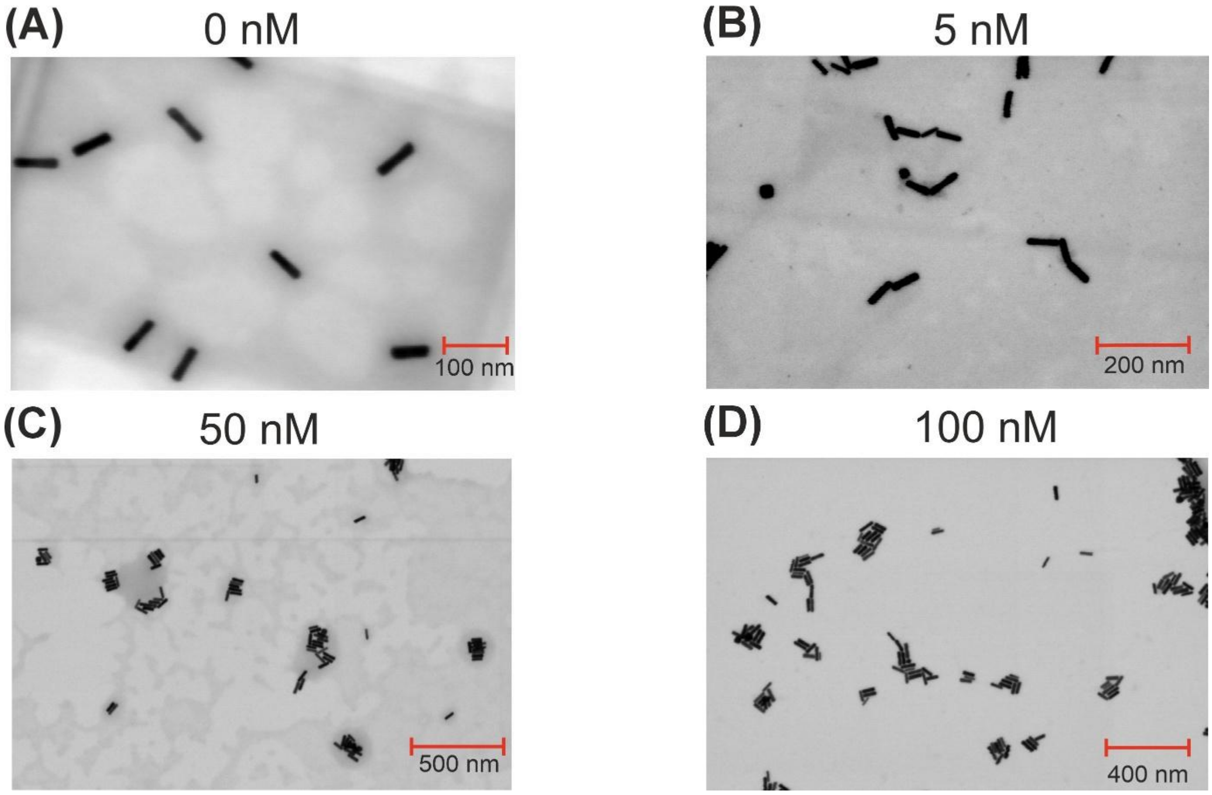


**Figure S8:** STEM images of target DNA-mediated AuNR assemblies formed at different target DNA concentrations: (A) 0 nM, (B) 5 nM, (C) 50 nM, and (D) 100 nM. In the absence of target DNA, the AuNRs remain predominantly dispersed. Upon addition of target DNA, assembly formation is observed through the appearance of dimers, short chains, and larger multi-particle clusters. Increasing the target DNA concentration leads to a progressive increase in the size and population of AuNR assemblies, consistent with the concentration-dependent optical response observed in the UV–Vis absorption measurements.

### 10. Normal Raman spectra of nucleotide monophosphates

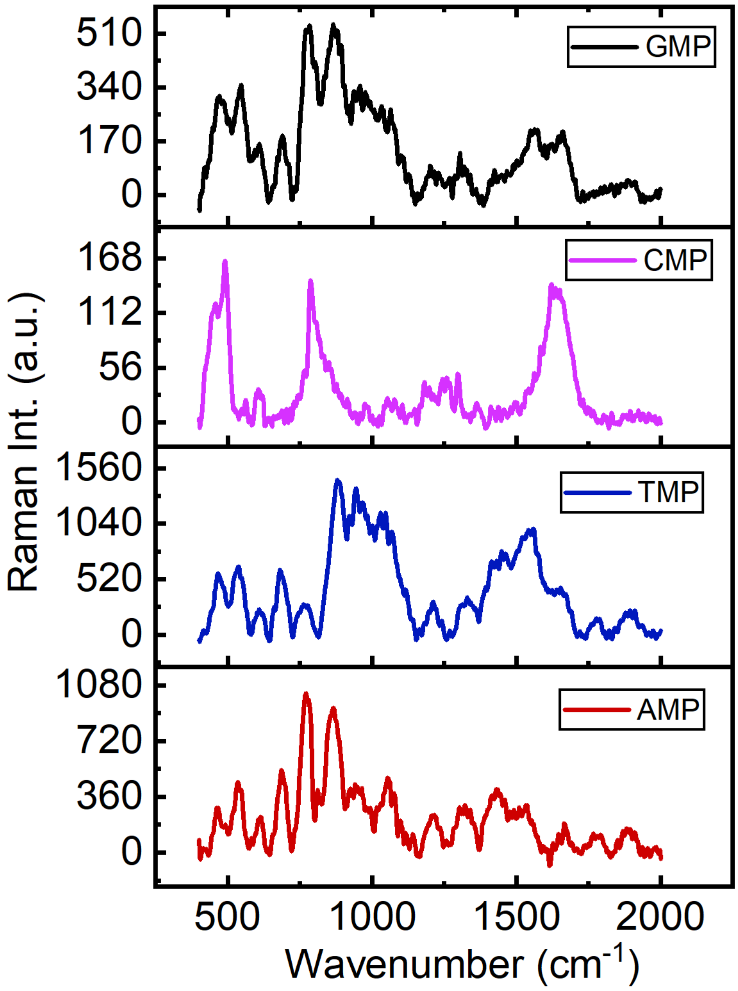


**Figure S9:** Normal Raman spectra of nucleotide monophosphates. Raman spectra of guanosine monophosphate (GMP, black), cytidine monophosphate (CMP, magenta), thymidine monophosphate (TMP, blue), and adenosine monophosphate (AMP, red) recorded at a concentration of 10 mM. The spectra were acquired under identical experimental conditions and are presented to facilitate assignment of the Raman bands observed in the SERS spectra of the target DNA. Among the four nucleotide monophosphates, AMP exhibits a prominent Raman feature in the 770–800 $cm^{-1}$ region, which is consistent with the characteristic band observed at approximately 773 $cm^{-1}$ in the SERS measurements.

**11. Normal Raman spectrum of the target DNA**

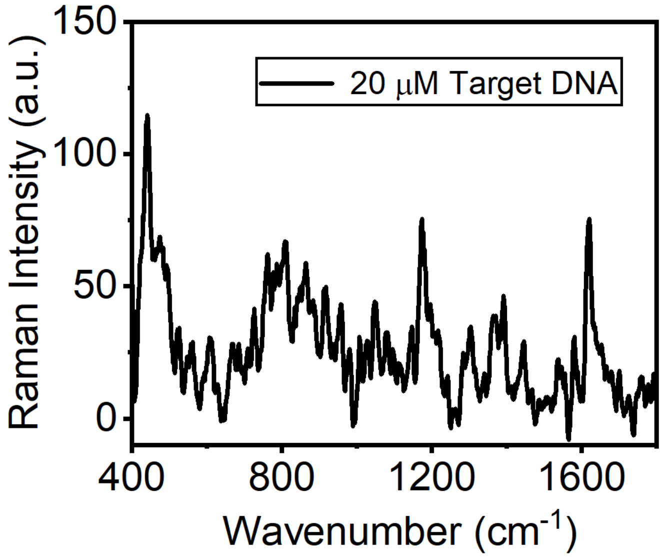


**Figure S10.** Normal Raman spectrum of the target DNA recorded at a concentration of 20 μM under identical experimental conditions used for the SERS measurements. The spectrum exhibits weak Raman features, highlighting the limited sensitivity of conventional Raman spectroscopy toward the target DNA in the absence of plasmonic enhancement.
